# Multiplex enCas12a screens show functional buffering by paralogs is systematically absent from genome-wide CRISPR/Cas9 knockout screens

**DOI:** 10.1101/2020.05.18.102764

**Authors:** Merve Dede, Megan McLaughlin, Eiru Kim, Traver Hart

## Abstract

Major efforts on pooled library CRISPR knockout screening across hundreds of cell lines have identified genes whose disruption leads to fitness defects, a critical step in identifying candidate cancer targets. However, the number of essential genes detected from these monogenic knockout screens are very low compared to the number of constitutively expressed genes in a cell, raising the question of why there are so few essential genes. Through a systematic analysis of screen data in cancer cell lines generated by the Cancer Dependency Map, we observed that half of all constitutively-expressed genes are never hits in any CRISPR screen, and that these never-essentials are highly enriched for paralogs. We investigated paralog buffering through systematic dual-gene CRISPR knockout screening by testing algorithmically defined ~400 candidate paralog pairs with the enCas12a multiplex knockout system in three cell lines. We observed 24 synthetic lethal paralog pairs which have escaped detection by monogenic knockout screens at stringent thresholds. Nineteen of 24 (79%) synthetic lethal interactions were present in at least two out of three cell lines and 14 of 24 (58%) were present in all three cell lines tested, including alternate subunits of stable protein complexes as well as functionally redundant enzymes. Together these observations strongly suggest that paralogs represent a targetable set of genetic dependencies that are systematically under-represented among cell-essential genes due to genetic buffering in monogenic CRISPR-based mammalian functional genomics approaches.

## Introduction

The adaptation of CRISPR-Cas9 system to genome-wide knockout screens in mammalian cells has greatly transformed the search for cancer specific genomic vulnerabilities that can be targeted therapeutically. Monogenic pooled library CRISPR-Cas9 knockout screens revealed that mammalian cells have as much as 3-4 times more essential genes than the previous RNAi technology was able to detect at the same false discovery rate (Hart et al., 2014). Moreover through immense monogenic screening efforts, multiple groups revealed lists of ~2000 highly concordant human essential genes, and comparison of CRISPR technology to orthogonal techniques such as random insertion of gene traps also showed consistent results (Blomen et al., 2015; Hart et al., 2015; Wang et al., 2015).

However, even with the CRISPR technology, the number of essential genes detected through these screens is still far less than the number of genes constitutively expressed in a given cell line. This phenomenon was previously observed in systematic gene knockout studies in *S. cerevisiae* (Giaever et al., 2002; Winzeler et al., 1999), where only 17% of yeast genes were essential for growth in rich medium (Winzeler et al., 1999). A closer look at the biological characteristics that define essentiality revealed a modular nature of gene essentiality (Hart et al., 2007) in which essentiality is not a characteristic of the protein or gene itself, but is rather defined by the protein complex to which the protein belongs. While genes that encode for members of a protein complex were shown to be more likely to be essential, paralogous genes were less likely to be essential (Gu et al., 2003). However, a later study showed that a binary classification of genes into essential and non-essential was misleading due to the context-dependent nature of gene essentiality and that 97% of yeast genes showed some growth phenotype under different environmental conditions (Hillenmeyer et al., 2008). A similar study in *C. elegans* (Ramani et al., 2012) suggested that virtually every gene is required for optimal growth in some condition.

Paralogous genes arise from gene duplication events, which is a mechanism to create new genes. While gene duplication can result in two functionally distinct genes over time, more frequently, the genes preserve a proportion of functional overlap through the process of subfunctionalization (Brookfield, 1997; Conant and Wolfe, 2008). In yeast gene deletion studies, singletons, which are genes without paralogs, were more than twice as likely as paralogous genes to be essential (Gu et al., 2003), indicating the role of paralogs in genetic buffering and suggesting that that paralogs can affect how the yeast cells can respond to genetic perturbations. The buffering ability of paralogs to each other’s loss can be explained by their functional redundancy. Double deletion studies of paralog gene pairs in yeast revealed that synthetic lethality occurred with depletion of both paralog pairs, resulting in a fitness defect that was more than the expected additive effect of individual gene depletions (DeLuna et al., 2008). Further analyses determined sequence similarity of paralog pairs as a predictive characteristic for the level of functional redundancy (Li et al., 2010). A major open question remains whether these findings hold true for human cells generally and cancer cells specifically.

Recent studies investigated paralog dependencies in monogenic genome-wide CRISPR-Cas9 knockout screens in human cells, revealing differential effects of paralogs on cellular fitness. One study showed that paralogs are less likely to be essential in whole-genome CRISPR knockout fitness screens than singleton genes (De Kegel and Ryan, 2019), while another study demonstrated that paralogs that form heterodimers are more deleterious to the cell compared to non-heterodimer forming paralogs (Dandage and Landry, 2019). However, these studies did not take into account the effect of tissue-specific expression of the paralog pairs.

In this study, using publicly available genome wide screen data of genetically heterogeneous cell lines from the Cancer Dependency Map initiative (Meyers et al., 2017; Tsherniak et al., 2017) we investigate paralogs among constitutively expressed never-essential genes as a set of targetable genetic dependencies that are systematically excluded in monogenic CRISPR-Cas9 knockout screening. We further demonstrate experimentally, using CRISPR/enCas12a multiplex knockouts, that dual-gene screens reveal synthetic lethality among targeted paralogs.

## Results

As part of our ongoing effort to understand differential gene essentiality, we looked at the relationship between gene expression and gene essentiality across hundreds of cancer cell lines. We looked at gene expression from all cell lines in the Cancer Cell Line Encyclopedia (CCLE) (Barretina et al., 2012), and considered the role of tissue-specific vs. constitutive gene expression. We took the mean and standard deviation of gene expression across 684 cell lines with high-quality CRISPR screens from the Avana 19Q4 data release (Meyers et al., 2017) and modeled the joint distribution with a linear combination of 2-d Gaussian mixture models. We find that three elements correspond to the three major populations in the data: constitutively expressed genes (high expression, low variance), never-expressed genes (low expression, low variance), and genes that show tissue-specific gene expression (“sometimes expressed” genes, high variance) (Figure 1A).

**Figure 1.**
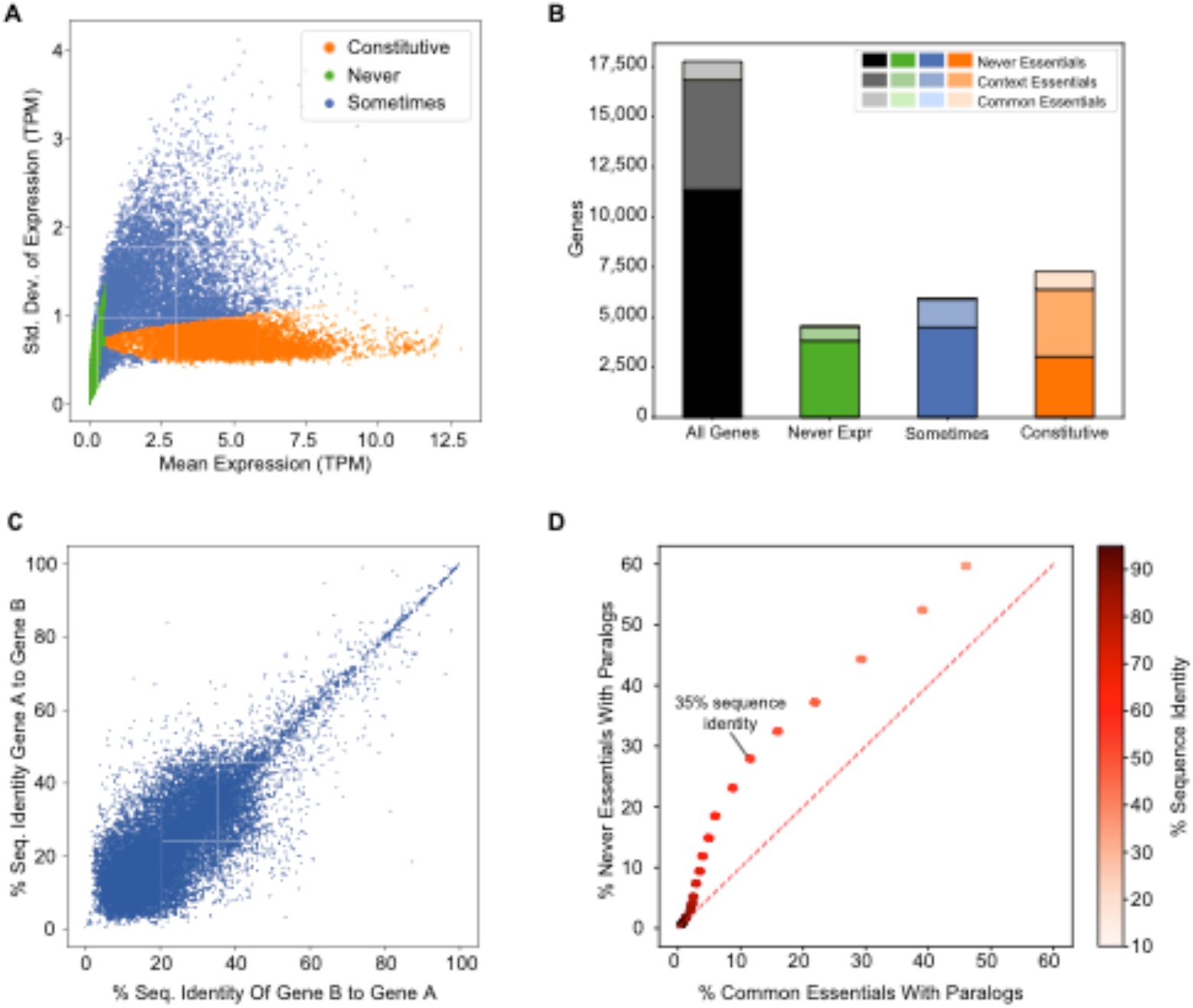
Paralogs are under-represented in CRISPR-Cas9 screens. A) Scatter plot of mean vs. standard deviation of log(TPM) gene expression in CCLE. Color coding by component of a 3-element Gaussian mixture model. (B) Fraction of context and common essentials as a function of gene expression (C) Paralog pairwise sequence identity among constitutively expressed paralogs. (D) Fraction of common essentials with a paralog vs. fraction of never essentials with a paralog, by paralog similarity.

We evaluated the fraction of essential genes in each population. Common essential genes are largely constitutively expressed, as expected, while context-dependent essential genes are divided across the constitutive expression and tissue-specific expression. Interestingly, among constitutively expressed genes, many are never essential in any CRISPR knockout fitness screen (3,032 of 7,282; 42%; Figure 1B). These observations regarding the constitutively expressed genes raised the question about why we observe so few essential genes in these genetically heterogenous screens. Based on work in yeast and nematodes (Ramani et al., 2012), we naively assumed that all constitutively expressed genes should be essential in some context, and hypothesized that some combination of environmental or genetic buffering masks the fitness consequences of individual gene knockouts.

It has been previously observed that paralogs are less likely to be essential in whole-genome CRISPR knockout fitness screens than singletons (De Kegel and Ryan, 2019), but this study did not correct for tissue-specific gene expression. To examine the role of paralog genetic buffering among constitutively expressed genes, we obtained the list of the paralogs of human protein coding genes from Ensembl Biomart (Zerbino et al., 2017) along with protein sequence similarity information (see Methods). After filtering for constitutively expressed genes, we observed that paralogs show a wide range of amino acid sequence similarity, with the majority showing relatively low identity (Figure 1C). To evaluate whether paralogs are enriched in constitutively expressed never-essentials (hereafter “never-essentials”), we adopted a sliding scale of sequence identity and measured, at each threshold, the fraction of never-essentials and the fraction of common essentials captured. As shown in Figure 1D, as sequence similarity stringency is relaxed, never-essentials are more likely to have a paralog than common essentials. At 35% or greater sequence similarity, nearly a third (27.9%) of constitutively expressed never-essentials have a paralog, compared with only 11.6% of common essentials.

To identify functionally redundant paralogs, we explored the Avana and Sanger data to find cases where loss of function of one member of a paralog pair resulted in increased dependency on the other (Figure 2A). We limited the search for functional redundancy to genes classified as constitutively expressed according to our model, which excludes false associations arising from tissue-specific expression of paralog family members. The search is further constrained by requiring that one member of the pair show loss of function, either through predicted deleterious mutation or by strong decrease in gene expression (see Methods), in a sufficient number of cell lines to result in a statistically significant difference in gene essentiality of the other member. By applying this test to 628 gene pairs in the Avana data and 295 gene pairs in Project Score (Figure 2A, Supp Table 1), we detected a total of 66 buffering interactions at a P-value < 0.01, of which 32 (48%) are common between the two sets (Figure 2B,C). Two well-described cases in the BAF (mammalian SWI/SNF) complex were immediately apparent: mutations in *SMARCA4* are strongly associated with dependency on paralog *SMARCA2* (P<10^−10^; Figure 2D), and mutations in *ARID1A* are associated with *ARID1B* dependency (P<10^−9^; Figure 2E). Expanding loss-of-function to include significantly depleted gene expression also reveals an emergent dependency on *RPP25L* when *RPP25* is depleted (P<10^−52^; Figure 2F). The two genes encode redundant subunits of RNAse P, a ribonuclease critical for maturation of tRNA, whose functional buffering was previously observed (Wang et al., 2015). A fourth example is *FAM50A/FAM50B* putative functional redundancy (Figure 2G). Interestingly, virtually nothing is known about the biological role of these genes. Unfortunately, the cell lines screened by CRISPR knockout libraries only contain LOF alleles of a fraction of the candidate paralogs, limiting this discovery avenue to a few dozen pairs.

**Figure 2.**
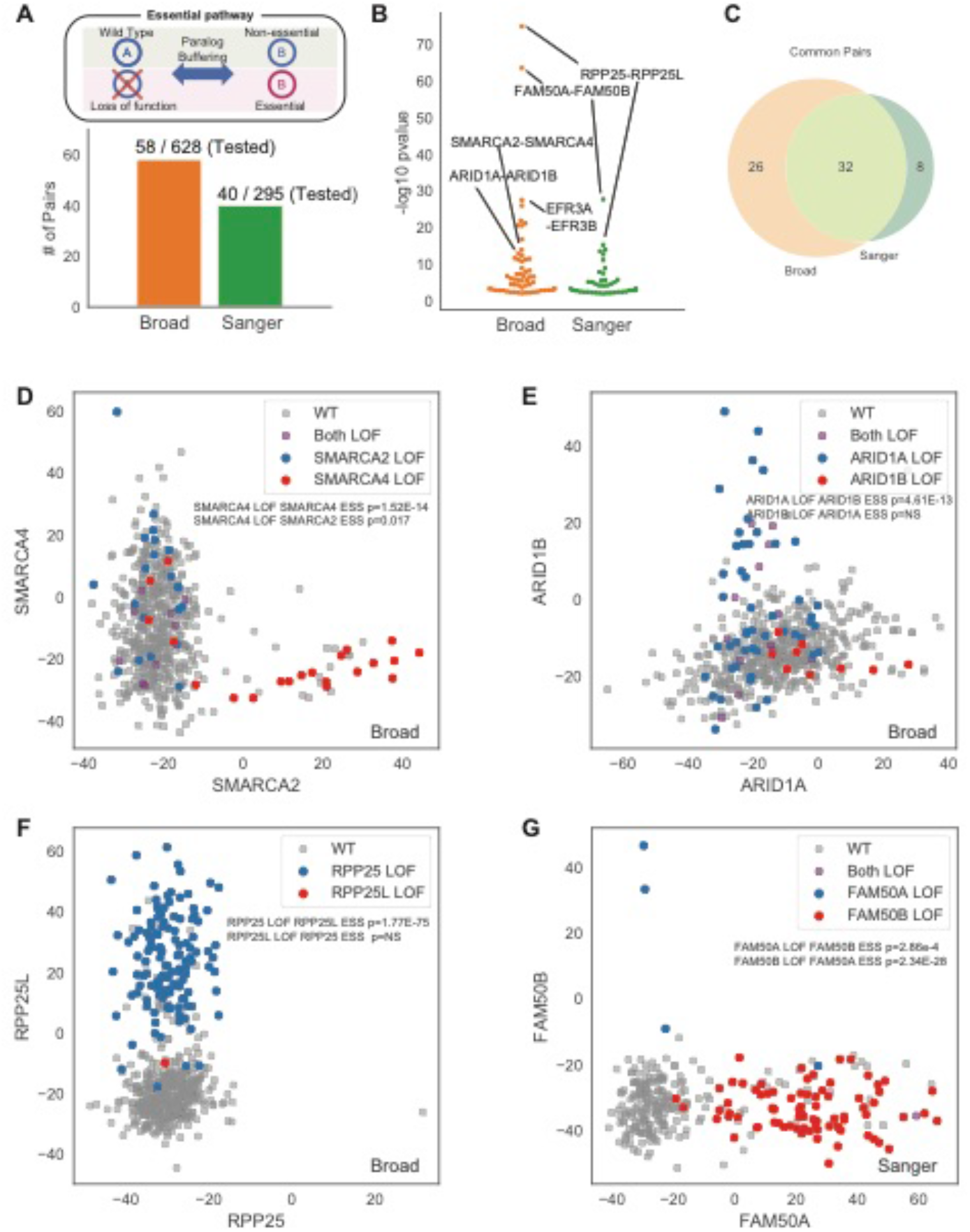
Computational detection of constitutively-expressed paralog buffering. (a) Method overview: loss of function in one paralog gives rise to gene essentiality in the other. Chart indicates number of pairs testable by this method. (B) Summary of results by dataset. (C) Overlap between Sanger (Project Score) and Broad (DepMap) computationally-derived synthetic lethals. (D-G) Scatter plots of Bayes Factors of paralog pairs, with loss of function (LOF) labeled.

Given the limitations of this computational approach, we sought to expand our knowledge of paralog buffering through systematic dual-gene CRISPR knockout screening. Cas12a, formerly Cpf1, offers an endogenous RNA endonuclease function that enables processing and utilization of multiple gRNA from a single polycistronic transcript (Zetsche et al., 2015) and the modified enCas12a enzyme offers superior performance in genetic screens in mammalian cells (Kleinstiver et al., 2019; Sanson et al., 2019). A key advantage of this system is that specific guide pairs can be synthesized in a single oligo, allowing one-step library design, a major advantage over multiplex Cas9 systems (Cong et al., 2013; Kabadi et al., 2014; Chen et al., 2015; Shen et al., 2017). We therefore sought to apply the enCas12a multiplex knockout system to systematically identify paralog synthetic lethals. In our hands, cells with enCas12a effectively knocked out EGFP (Supp Fig 1a) and achieved ~80% double knockout in a dual-guide construct targeting two cell surface markers (Supp Fig 1b).

We designed a dual-guide library targeting ~400 candidate paralog pairs. Gene pairs were selected based on several criteria, including amino acid sequence similarity, mRNA expression and co-expression, and whether either gene is frequently essential in DepMap. We manually five additional candidate gene pairs from the literature: *SMARCA2-SMARCA4*, *CHD1-CHD3*, *ME2-ME3*, *BCL2L1-MCL1*, and *BRCA1-PARP1*, for a total of 403 targeted gene pairs. For each gene, three CRISPR RNA (crRNA) were selected using the DeepCpf1 scoring scheme (Kim et al., 2018). Each gene pair was targeted with all 9 combinations of guides, in both A-B and B-A orientations, for a total of 18 clones targeting each pair, and we paired gene-targeted crRNA with guides targeting known nonessential genes to evaluate single-knockout phenotype (Figure 3A). We additionally targeted 50 essential and 50 nonessential genes as quality controls for the screens.

**Figure 3.**
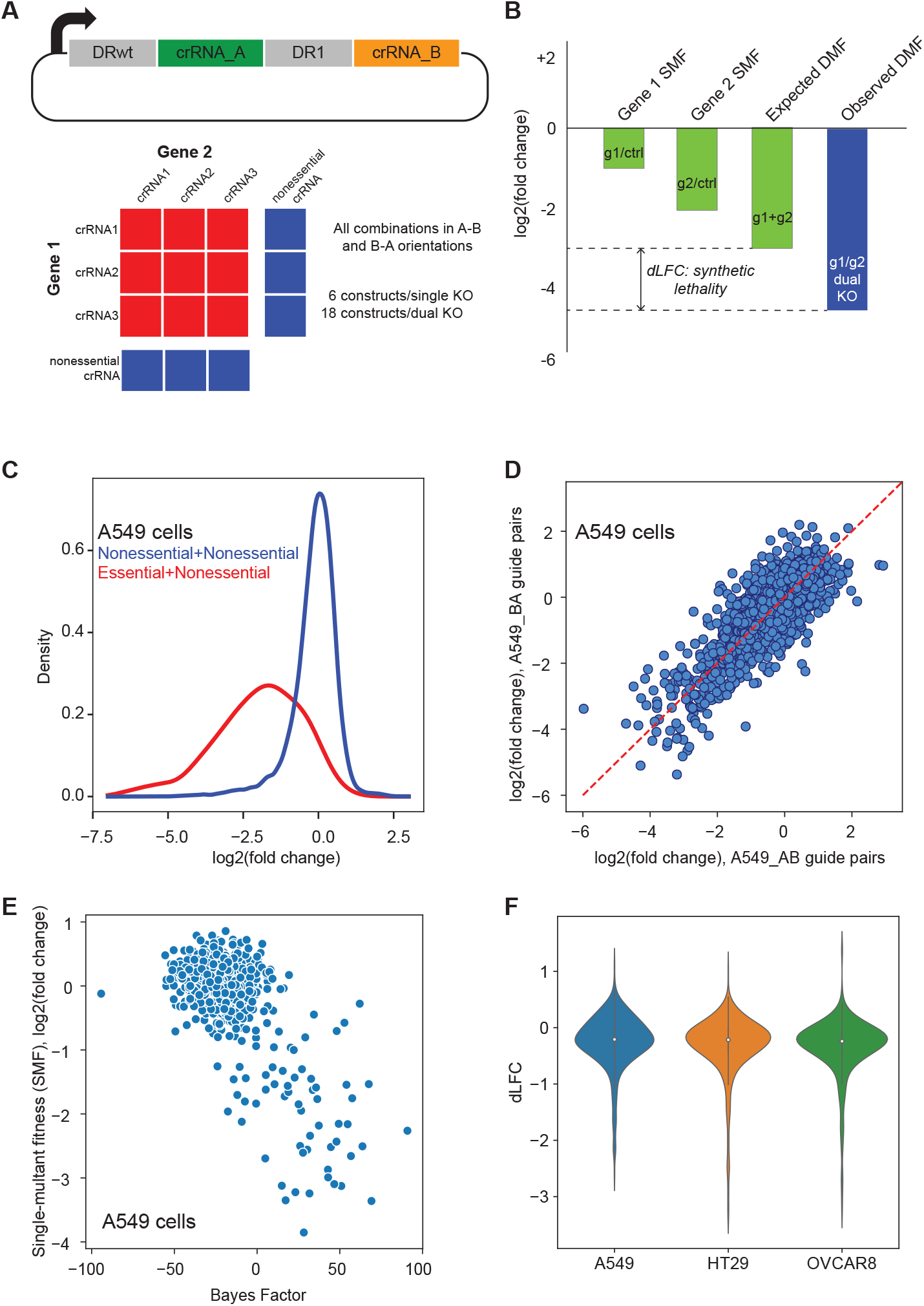
Multiplex gene knockout with enCas12a. (A) Experimental design. EnCas12a crRNA dual-guide array design. Each construct targets two genes; each gene is targeted by 3 crRNA; each candidate paralog gene pair is targeted by 18 gene-gene constructs, with six gene-control constructs per gene (including both A-B and B-A orientations). (B) Evaluating synthetic lethality. Single mutant fitness (SMF) is the mean log fold change of control guides targeting a single gene. Expected double mutant fitness (DMF) is the sum of SMF. Observed DMF is the mean log fold change of dual-targeting constructs. Delta log fold change (dLFC) is the difference between observed and expected fold change. (C) QC plot showing separation of SMF of constructs targeting control essential and nonessential genes. (D) Scatter plot of all mirror constructs (same crRNA in A-B and B-A orientations) showing lack of positional effects. (D) SMF in this screen vs. gene BF in Avana data. (E) Distribution of dLFC scores for 403 gene pairs in each cell line.

We transduced the library into enCas12a-expressing cells from three cell lines: A549, a KRAS-driven lung cancer cell line; HT29, a BRAF-mutant colorectal cancer cell line, and OVCAR8 ovarian cancer cells. Cells were passaged in three replicates for 10 doublings and the relative abundance of each dual-guide construct was measured by 75-base single end sequencing of the target amplicon, with fold changes measured relative to abundance in the plasmid pool. Quality control steps including abundance and distribution of read counts, clustering of raw read counts and fold changes, and separation of essential and nonessential control genes indicated effective screen performance (Figure 3C and Supp Fig 1). Additionally, high correlation of A-B and B-A guide pairs (Figure 3D) indicate negligible positional bias in the enCas12a guide arrays. We therefore included both A-B and B-A pairs in all subsequent fitness calculations.

To calculate genetic interaction/synthetic lethality, we measured the single mutant fitness (SMF) for each gene as the mean fold change of the gene-control constructs. For control essential genes, SMF in our enCas12a screen correlates with BAGEL-derived Bayes Factor scores for the DepMap screens in the same cell lines (Figure 3E). We then calculated the observed double mutant fitness (DMF) as the mean log fold change of the dual-gene knockout constructs (18 constructs per gene pair), and compared it to the expected DMF, the sum (in log space) of each gene’s SMF (Figure 3B). As has been widely observed in genetic interaction screens, most digenic knockouts do not result in an unexpected phenotype; here we observe that the distribution of “delta log fold change” (dLFC) values has most of its mass around zero (no synthetic effect), with a long tail of negative (synthetic sick/lethal) dLFC scores (Figure 3F).

To compare across screens, we converted dLFC scores to a robust Z score, zdLFC, by truncating the top and bottom 2.5% of dLFC scores (Supp Fig 1, Supp Table 2, Supp Table 3). At a zdLFC score < −3, all three screens showed high concordance, with 19 of 24 (79%) synthetic lethals present in at least two out of three cell lines and 14 of 24 (58%) present in all three (Figure 4A-B). Many top-scoring hits show strong concordance with other data corroborating a functional buffering/synthetic lethal relationship. RNA helicases *DDX19A* and *DDX19B* show characteristics synthetic lethality as described by De Kegel and Ryan (De Kegel and Ryan, 2019) across DepMap cell lines, DDX19A is strongly essential only when DDX19B is expressed at low levels (Figure 4C). Similarly, TIAL1 low expression is associated with TIA1 increased essentiality. Synthetic lethality between these genes is corroborated by a dual-gene knockout screen using a hybrid Cas12/Cas9 system (Gonatopoulos-Pournatzis et al., 2020). Genes *CNOT7* and *CNOT8* encode alternate subunits of the CCR4-NOT complex, a critical regulator of eukaryotic gene expression (Lau et al., 2009). Other subunits are sporadically essential in our three cell lines (Figure 4D) but frequently essential across DepMap data (Lenoir et al., 2018; Meyers et al., 2017; Tsherniak et al., 2017), consistent with a constitutively essential protein complex. Moreover, CNOT7 essentiality is weakly but significantly anticorrelated with CNOT8 mRNA expression (Pearson correlation coefficient −0.21, P<10^−6^). Likewise, *COPS7A* and *COPS7B* encode alternate, replaceable subunits of the COP9 signalosome complex; other subunits are irreplaceable and are uniformly essential in these cell lines (Figure 4E).

**Figure 4.**
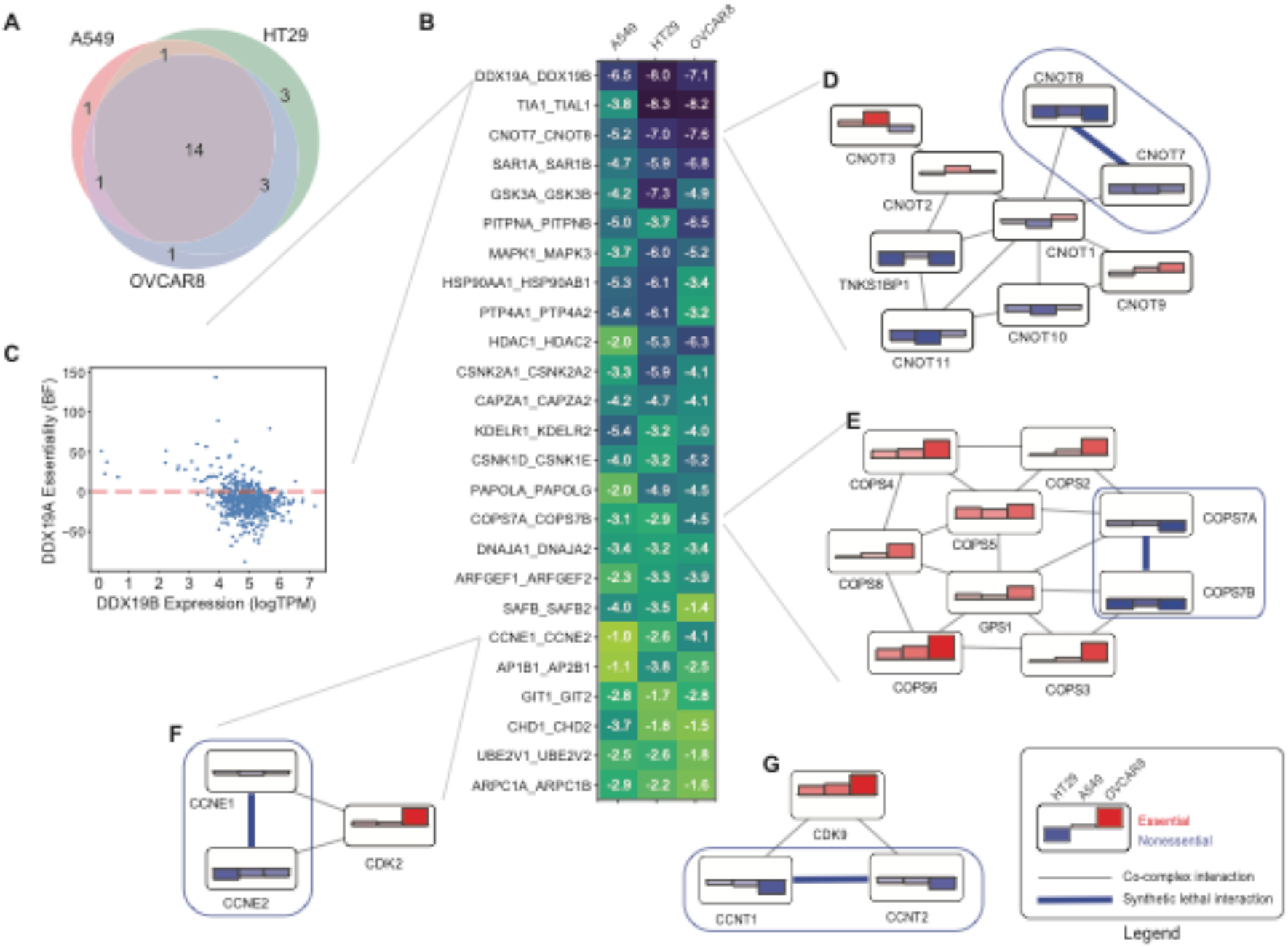
Synthetic lethal paralogs. (A) Overlap of 24 hits with zdLFC < −3 in any of the three cell lines. (B) Heatmap showing zdLFC score for top 25 hits. (C) *DDX19A* gene essentiality (BF) vs. *DDX19B* gene expression (logTPM) in Avana data. (D) Selected co-complex interactions in CCR4-NOT complex shows gene essentiality of several subunits and synthetic lethality of nonessential, alternate subunits *CNOT7/CNOT8*. (E) Co-complex interactions of COPS9 signasosome complex, showing strong gene essentiality of required components and synthetic lethality of alternate subunits *COPS7A/COPS7B*. (F) Synthetic lethality of *CCNE1/CCNE2* in OVCAR8 cells, where *CDK2* is highly essential. (G) Synthetic lethality of *CCNT1/CCNT2*, not shown in B, and essentiality of *CDK9*.

Importantly, we note that synthetic lethality, even among paralogs, can also be context-dependent. Cyclin paralogs are often redundant interaction partners with their cognate cyclin-dependent kinases; here, CCNE1 and CCNE2 are synthetic lethal where CDK2 is highly essential, especially in OVCAR8 (Figure 4F). Similarly, CCNT1-CCNT2 show weaker but significant synthetic lethality (zdLFC < −2.5 in A549 and < −1 in the other two cell lines) while their binding partner, CDK9, is highly essential in all three (Figure 4G). Though the synthetic lethal relationships between SWI/SNF complex members *ARID1A/ARID1B* and *SMARCA2/SMARCA4* are well described in the literature and are detected in large scale screening data, their synthetic lethality only occurs where the SWI/SNF complex is itself essential. We test four paralog pairs in the BAF complex: *ARID1A/ARID1B, SMARCA2/SMARCA4, SMARCC1/SMARCC2,* and *SMARCD1/SMARCD2*, but we detect no synthetic lethal interactions, most likely because the complex itself is not essential in the cell lines we tested.

## Discussion

CRISPR technology has revolutionized mammalian functional genomics and cancer targeting by leveraging endogenous DNA repair machinery to generate gene knockouts on a genomic scale. Extensive screening of cancer cell lines has been performed under the DepMap and Project Score initiatives to identify context-specific weaknesses and cancer biomarkers. Analyses of this data have revealed activation of oncogenic pathways and oncogene dependencies (Tsherniak et al., 2017) as well as biomarker type dependencies such as Werner helicase, WRN, in colorectal and ovarian cell lines with MSI (Behan et al., 2019; Chan et al., 2019). However, despite these efforts, questions about what might be systematically missing from these data have, to our knowledge, not been rigorously explored.

We note that there are about 7,000 genes that are constitutively expressed in each cell, but only about half of these are ever detected as essential. Studies in model organisms suggest that virtually every gene shows a growth phenotype under some environmental condition (Hillenmeyer et al., 2008; Ramani et al., 2012). It is unknown whether this holds true for individual mammalian cells, though tumors are often modeled as though they are colonies of single-celled organisms. It is also the case that most genetic screens of tumor cells are carried out under permissive growth conditions, minimizing nutrient and oxidative stress to maximize growth rate and improve detection of dropouts. Thus, the degree of environmental buffering is largely unknown for these constitutively expressed never-essentials.

However, these never-essentials are highly enriched for paralogs. They are ~3 times more likely to have a paralog than always-essentials, suggesting that functional redundancy by related genes masks detection of a substantial population of genes in monogenic CRISPR knockout screens. This has profound implications for efforts to match targeted drugs with tumor genotypes, and to discover new candidate drug targets. Targeted small molecules often don’t discriminate, or discriminate poorly, between closely related paralogs, and it is often their promiscuity rather than their specificity that renders them effective. For example, MEK inhibitor trametinib effectively targets the protein products of both *MAP2K1* and *MAP2K2*, redundant kinases downstream of RAS/RAF oncogenes, but the functional redundancy of these genes renders them both invisible to monogenic CRISPR screens, even in RAS/RAF backgrounds (Kim et al., 2019).

Recent developments in CRISPR screening technology enable effective genetic targeting of multiple genes simultaneously. Cas12a, previously known as Cpf1, is able to process a polycistronic mRNA to generate multiple CRISPR RNAs (crRNAs). This makes multiplexing much easier compared to inefficient Cas9 based multiplex systems which requires each guide RNA to be expressed by its own promoter. The improved version of this enzyme, enCas12a (Kleinstiver et al., 2019), coupled with an effective guide design algorithm (Kim et al., 2018) presents a powerful platform for multiplex genetic perturbation. Multiplex guide libraries can be synthesized directly, without requiring additional targeted or random mixing cloning steps, allowing direct assay of specific gene pairs as described here with roughly the same level of effort as a now-standard Cas9 monogenic screen. The robustness of predicted guide cutting efficiency remains untested, given the relatively small amount of enCas12a data available, suggesting adopters of this technology should err toward caution when deciding on parameters for new experiments (e.g. number of guides per gene, number of gene-vs-control guide pairs). Nevertheless, as we demonstrate here, this platform holds enormous potential for exploring the stability and plasticity of genetic interactions in human cells.

## Methods

### Essentiality data generation

A raw read count file of CRISPR pooled library screens for 690 cell lines using Avana library (Meyers et al., 2017) (Broad DepMap project 19Q4) was downloaded from the data depository (https://depmap.org/portal/). Also, we downloaded Project Score (Sanger) screen (Behan et al., 2019) raw read counts for 323 cancer cells from the data depository (https://score.depmap.sanger.ac.uk/). We filtered the dataset to keep only the protein-coding genes for further analysis and updated their names using HGNC (DeLuna et al., 2008) and CCDS (Farrell et al., 2014) database. We discarded sgRNAs targeting multiple genes in Avana library to avoid genetic interaction effects. The raw read counts were processed with the CRISPRcleanR (Iorio et al., 2018) algorithm to correct for gene-independent fitness effects and calculate fold change. After that, the CRISPRcleanR processed fold changes of each cell line were analyzed through updated BAGEL2 build 114 (https://github.com/hart-lab/bagel). In comparison with published BAGEL version v0.92 (Hart and Moffat, 2016), the updated version employed a linear regression model to interpolate outliers and 10-fold cross validation for data sampling. Essentiality of genes was measured as Bayes Factor (BF) based on gold standard reference sets of 681 core essential genes and 927 nonessential genes (Hart et al., 2014; Hart et al., 2017). Positive BF indicates essential genes and negative BF indicates non-essential genes. Lists of core essential genes and nonessential genes used in this study have been uploaded on the same repository with BAGEL2 software. To correct unexpected essentiality by sgRNAs targeting non-protein coding regions in addition to desired target protein coding gene, the multi-targeting effect of sgRNAs has been corrected using BAGEL2 -m option. The screen quality was evaluated by using “precision-recall” function in BAGEL2 software, and F-measure (BF = 5), which is the harmonic mean of precision and recall at BF 5, was calculated for each screen. Finally, 581 cell lines for Broad screen and 320 cells for Sanger screen were selected for further study by F-measure threshold 0.8 to prevent noise from marginal quality of screens.

### Defining constitutively expressed genes with GMM modeling

We utilized the log2 transformed RNA-seq TPM expression data from DepMap Data Portal expression data for Avana19Q4 release for 684 cell lines (Meyers et al., 2017; Tsherniak et al., 2017). The standard deviation of expression versus the mean expression values for all genes assayed in the Avana library (N=17,755) across all cell lines, for which the expression data was available, were plotted. Python 3.6.9 package sklearn and its GaussianMixture function was used to classify genes into 3 groups based on mean and standard deviation of mRNA expression. The group with the least expression and low standard deviation was labeled as never expressed, the second group with very high standard deviation and a range of mean expression values was labelled as sometimes expressed and the constitutively expressed group with high mean expression and low standard deviation was classified as constitutively expressed genes. With this classification, we identified 7,282 always expressed, 4,544 never expressed and 5,929 sometimes expressed genes in the Avana dataset.

### Paralogs

The human paralogous gene pairs for the protein coding genes were utilized from Ensemble Release 95 Biomart with GRCh38.p12 genome assembly (Zerbino et al., 2017). This release of Ensemble estimates paralogues from gene trees that are constructed with HMM as described in more detail at http://www.ensembl.org/info/genome/compara/homology_method.html. Other information such as chromosome location, paralogue percent sequence identity to human target gene and percent sequence identity of target gene to the paralogous gene were also downloaded. After removing duplicate gene pairs and filtering for constitutively expressed genes, the percent sequence identities of all human target genes to their paralogs were plotted against the percent sequence identities of the paralogs to the target human gene to reveal that the majority of the human paralogous gene pairs had low percentage sequence similarity. The paralog pairs which were both in always expressed gene list were identified and were binned according to different thresholds for percent sequence identity from a range of 10-95%. For each bin, the percentage of constitutively expressed never essential genes with paralogs and the percentage of common essential genes (defined in (Dede et al., 2020)) with paralogs were calculated and their distributions were plotted. For downstream analysis, always expressed paralog pair lists were generated for each sequence identity threshold.

### Investigation of evidences of functional redundancy between paralog genes in CRISPR screens

To investigate evidence for the functional redundancy between paralog genes in Broad and Sanger screens, we tested whether a gene is essential when the other paralog partner suffers loss of function. Firstly, we defined loss of function (LOF) call combining damaging mutations calls (frameshift or nonsense) adopted from CCLE mutation data (Ghandi et al., 2019) depletion of expression (mean log TPM < 1.0, CCLE RNA-seq) or deletion (copy number < 0.1, CCLE Copy number data). Then, we conducted statistical test of synthetic essentiality which is defined that a gene is essential when the other paralog partner lost the function. One to one paralog pairs with at least 30% sequence similarity were considered for this analysis to maximize the number of paralog pairs. We considered only pairs whose genes have at least two LOF calls and are essential in at least two cell lines. P-value was calculated by the one-sided Fisher’s exact test on the 2×2 contingency table of the number of cells classified by LOF and essential (BF > 10). We addressed pairs bidirectional ways, which test a significance of essentiality of gene A upon LOF of gene B and *vice versa*. A total of 57 pairs among 628 tested pairs in the Broad dataset and 40 pairs among 295 tested pairs in the Sanger dataset passed a threshold of P-value < 0.01. Thirty-two pairs were common to both datasets.

### Identification of paralog test set

To identify and predict paralog pairs which we presumed to be enriched in synthetic lethal interactions, we used multiple parameters including percent sequence similarity, mean expression, standard deviation of expression, co-expression and gene essentiality profiles. We built a network of paralogous gene families using Cytoscape (Shannon, 2003) and filtered them initially for protein sequence similarity greater than or equal to 45%, mean expression (logTPM) >1.5, standard deviation of expression <1.25, co-expression Pearson correlation coefficient >0.1. Finally, we removed genes that were essential in more than 30 cell lines to obtain a set of 400 paralog pairs of interest.

### Vectors

The following vectors were a kind gift from John Doench:

pRDA_174 (enzyme expression): EF1a promoter drives EnCas12a enzyme expression; lentiviral vector; confers blasticidin resistance (Addgene #136476).
pRDA_052 (guide expression): U6 promoter drives gRNA expression; vector contains AsCas12a direct repeat upstream of dual BsmBI sites for insertion of guide arrays; lentiviral vector; confers puromycin resistance (Addgene #136474).
pRDA_221 (positive control): confers constitutive of short half-life EGFP and expression of two guides targeting EGFP; lentiviral vector; confers puromycin resistance (Sanson et al., 2019).
pDV204 (positive control): U6 promoter drives expression of guides targeting cell-surface markers CD47 and CD63; derived from pRDA_052 using guide sequences from Sanson et al., 2019.

### Library production

An oligonucleotide pool comprising 12k dual guide arrays was synthesized by CustomArray based on the following template: 5’*TATCTTGTGGAAAGGACGAAAC*ACCGG**TAATTTCTACTCTTGTAGAT**NNNNNNNNNNNNNNNNNNNNNNN<u>**TAATTTCTACTGTCGTAGAT**</u>nnnnnnnnnnnnnnnnnnnnnnnTTTTTTGAATT *CGCTAGCAAGCTTGGCGTAAC*-3’. The 145 nt fragment included the wildtype direct repeat for AsCas12a (bold) and an engineered variant direct repeat (bold underlined) (Sanson et al., 2019) for 23 nt guide sequences in the first (uppercase N) and second (lowercase n) positions, respectively. Flanking sequences (italic) enabled PCR amplification of the pool and cloning into BsmBI-linearized pRDA_052 by Gibson assembly.

The pool of guide arrays was amplified using Kapa HiFi 2X HotStart ReadyMix (Roche) using 10 ng of starting template per 50 μL reaction using primers DV202 (5’-TATCTTGTGGAAAGGACGAAAC) and DV203 (5’ GTTACGCCAAGCTTGCTAGCG) at 0.3 μM final concentration and the following conditions: initial denaturation at 95°C for 3 minutes, followed by twelve cycles of 20 s at 98°C, 20 s 60°C, 20 s at 72°C using a ramp rate of 2.0°C/second, final extension at 72°C for 5 minutes.

Full-length amplicon (145 bp) was purified by non-denaturing polyacrylamide gel electrophoresis using precast 10% acrylamide TBE gels (Bio-Rad). The guide expression vector pRDA_052 was digested with BsmBI (New England Biolabs), de-phosphorylated with Antarctic phosphatase (New England Biolabs) and concentrated using PCR cleanup columns (Life Technologies). Vector and insert were quantified using fluorimetry (Qubit dsDNA High Sensitivity kit, ThermoFisher). DNA assembly reactions using 0.4 pmol insert and 0.1 pmol vector per 20 μL HiFi Master Mix (New England Biolabs) were incubated at 50°C for 1 h, re-digested with BsmBI, and desalted (Monarch low volume elution columns, New England Biolabs) for electroporation into Endura electrocompetent cells (Lucigen). After 1 h recovery at 37°C, the bacteria were diluted 1:100 in 2xYT containing 200 μg mL^−1^ carbenicillin (AMS Bio) and grown at 30°C for 16 h. Transfection grade plasmid was purified (PureLink HiPure Maxiprep, Invitrogen) and its guide arrays were sequenced to confirm complete and uniform library representation.

### Cell Culture

A549 and HT29 cells were a kind gift from Tim Heffernan. OVCAR8 cells were a kind gift from Phil Lorenzi. Cell line identities were confirmed by STR fingerprinting by MD Anderson’s Cytogenetics and Cell Authentication Core (Powerplex 16 Locus High Sensitivity Assay, Promega).

Cells were grown at 37°C in humidified 5.0% CO_2_ atmosphere and passaged 2-3 times per week to maintain exponential growth. A549 and HT29 were grown in HEPES-modified DMEM (Sigma D7161); OVCAR8 was grown in HEPES-modified RPMI (Sigma R5886). Base media were supplemented with 10% FBS (Sigma), 1 mM sodium pyruvate (Gibco), 2 mM L-alanine-L-glutamine (Gibco), 1X penicillin-streptomycin (Gibco) and 100 μg mL^−1^ Normocin (Invivogen). Antibiotic-free cultures were routinely tested for mycoplasma contamination (PlasmoTest, Invivogen).

### Screens

Lentivirus was produced by the University of Michigan Vector Core. Virus stocks were not tittered in advance: all transductions were performed in multiple plates with a range of virus volumes and 8 μg mL^−1^ polybrene (EMD Millipore), but only the pool with the most optimal transduction efficiency was expanded and/or screened.

First, stable EnCas12a expression was engineered by transduction with pRDA_174 at low MOI (10% - 20% transduction efficiency). Non-transduced cells were eliminated by selection with 10 μg mL^−1^ blasticidin (Invivogen). Selection was maintained until non-transduced controls reached 0% viability twice in succession (~ ten doublings). Editing efficiency was confirmed by transduction with control vectors targeting EGFP (pRDA_221) or cell surface markers CD47 and CD63 (pDV204) and flow cytometry. Cell lines lacking EnCas12a expression served as controls. Conjugated fluorescent antibodies and isotype controls were from BioLegend.

Second, enzyme-expressing pools were transduced with guide array virus. Multiple sub-confluent 15 cm plates were transduced to achieve a minimum of 12M unique transductants without exceeding 50% transduction efficiency. Non-transduced cells were eliminated by 72 h treatment with 2 μg mL^−1^ puromycin (Gibco).

After puromycin selection was complete, three replicates were seeded, using 12M viable cells per replicate, i.e. ~1,000 cells per guide array. Screens were fed fresh medium every 2-3 days and passaged before reaching 80% confluency. Each replicate was re-seeded with 12M viable cells to maintain coverage. Remaining cells were stored in 30M aliquots at −80°C in cryopreservation medium (CellBanker 2, ZenoAq). Screens were terminated when replicates reached ten doublings.

### Sequencing

Genomic DNA (gDNA) purification was automated in 24-well plates using a Kingfisher Flex instrument (ThermoFisher) and magnetic bead-compatible reagents (Mag-Bind Blood and Tissue DNA HDQ, Omega Biotek). Purified gDNA was eluted in 10 mM Tris-HCl pH 8.0, 1 mM EDTA and quantified by fluorimetry (Qubit dsDNA Broad Range kit, ThermoFisher).

Illumina-compatible guide array amplicons were amplified from gDNA in one step, as described in Sanson et al., 2019). Indexed PCR primers were synthesized by Integrated DNA Technologies using Illumina’s standard 8nt indexes (D501-D508 and D701-D712). The forward primer design was 5’-**AATGATACGGCGACCACCGAGATCTACAC**NNNNNNNN**ACACTCTTTCCCTACACGACGCTCTTCCGATCT***CTTGTGGAAAGGACGAAACACCG (5’-* **i5 flow cell adapter** - i5 index - **i5 read1 primer binding site** - *U6 annealing sequence)*. The reverse primer design was 5’- **CAAGCAGAAGACGGCATACGAGAT**NNNNNNNN**GTGACTGGAGTTCAGACGTGTGCTCTTCCGATCT***GTTACGCCAAGCTTGCTAGCGAATTC* (*5’-* **i7 flow cell adapter** - i7 index - **i7 read2 primer binding site** - *pRDA_052 annealing sequence.)* Guide arrays were amplified from 80 μg of gDNA per replicate in multiple reactions, not exceeding 10 μg per 100 μl PCR volume. 80 μg represents at least 500 cells per guide array for these hypotriploid cell lines (www.ATCC.org).

Each 100 μL reaction contained 0.5 μM of each primer, 200 μM dNTPs and 1.25 μL of ExTaq polymerase (Takara). Guides were amplified using a slow ramp rate (2.0 °C/second) and minimum cycle number to limit bias, as follows: initial denaturation at 95°C for 60 s, followed by 28 cycles of 30 s at 94°C, 30 s at 52.5°C, 30 s at 72°C, final extension at 72°C for 10 m. *Please note that Sanson et al. (2019) now recommend using Titanium Taq plus DMSO (Takara) and we have observed slightly better mapping rates for Titanium Taq amplicons.* The ~200 bp indexed amplicons were purified by size selection (2% agarose, E-Gel SureSelect II, ThermoFisher), quantified (QuBit, ThermoFisher) and pooled. Sequencing was performed using custom read primer oligo1210 (5’-CTTGTGGAAAGGACGAAACACCGGTAATTTCTACTCTTGTAGAT) (HPLC purified, Integrated DNA Technologies) using NextSeq 1 × 75 nt High Output reagents (Illumina).

## Supplementary Figures

**Supplementary Figure 1:**
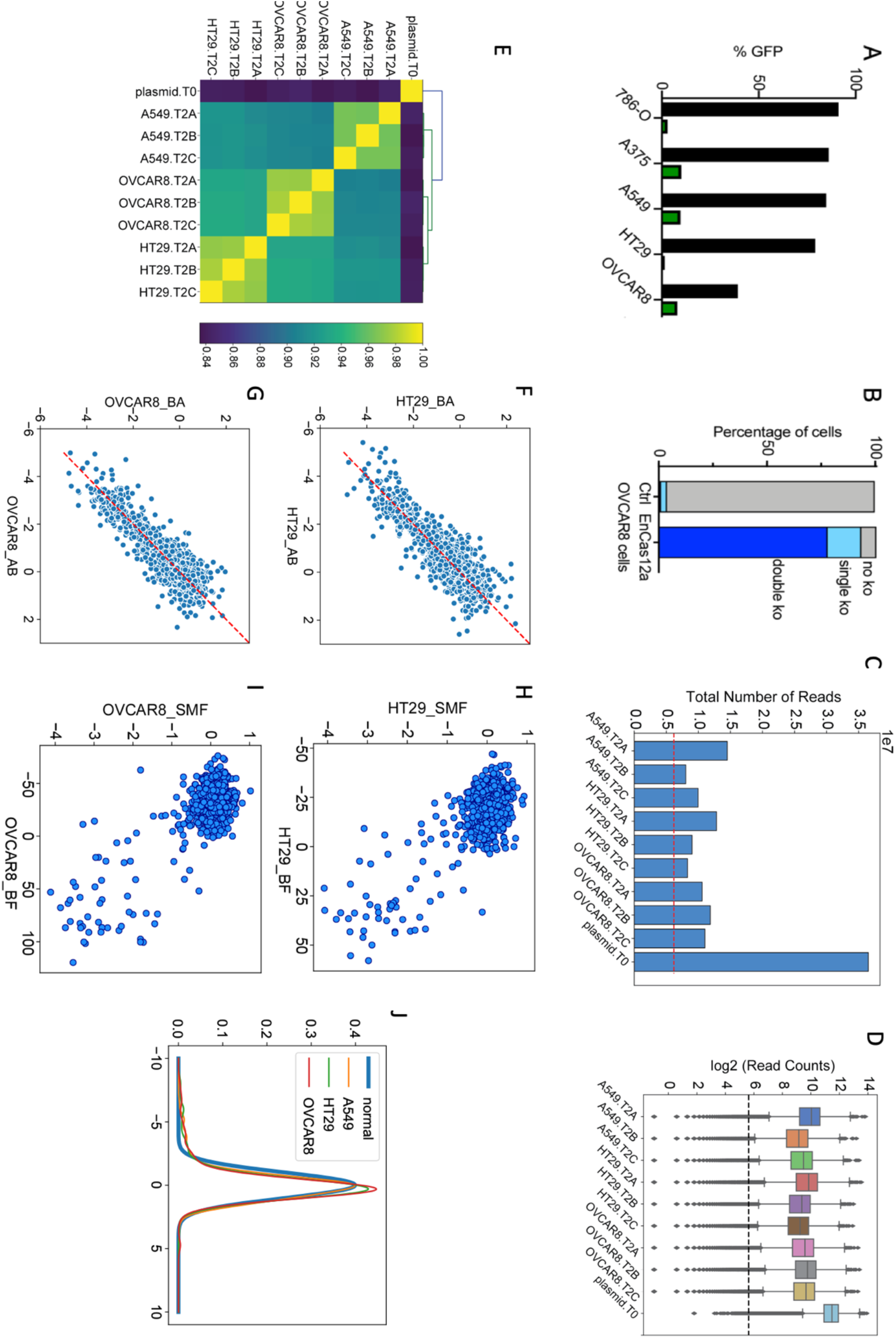
(A) Knockout of GPF with a crRNA targeting GFP in enCas12a knock-in cells. (B) Targeting two cell surface markers with a dual-guide crRNA in EnCas12a-expressing OVCAR8 cells. (C) Total amplicon reads for the paralog screen. Dashed line indicates 500x sequencing depth (6m reads for 12k library). (D) Distribution of reads (boxplot) indicates good library representation in each sample. (E) Clustering of normalized read counts consistent with high-quality screen data. (F, G) Lack of positional bias in mirror constructs containing the same two crRNA in A-B and B-A orientations. (H, I) SMF in this screen vs. BF from Avana data. (J) Z-transformation of distribution of dLFC (zdLFC) after truncating top/bottom 2.5% of values approximates a normal distribution.

## Supplementary Tables

Supplementary Table 1: Table of computational scores for predicted paralog pairs.

Supplementary Table 2: Table of zdLFC scores from the paralog screen.

Supplementary Table 3: Table of read counts of the paralog screen.

## Acknowledgments

Vectors pRDA_174 (Addgene #136476) and pRDA_052 (Addgene #136474) were a kind gift from John Doench. MD and MM were supported by the Cancer Prevention Research Institute of Texas (CPRIT) grant RR160032. TH is a CPRIT Scholar in Cancer Research, and is supported by NIGMS grant R35GM130119 and MD Anderson Cancer Center Support Grant P30 CA016672. TH is a consultant for Repare Therapeutics.

